# Alterations in Causal Functional Brain Networks in Alzheimer’s Disease: A resting-state fMRI study

**DOI:** 10.1101/2024.05.12.593795

**Authors:** Rahul Biswas, SuryaNarayana Sripada

## Abstract

**Background:** Alterations in functional connectivity (FC) of the brain is known to predate the onset of clinical symptoms of Alzheimer’s disease (AD) by several decades. Identifying the altered functional brain networks in AD can help in its prognosis and diagnosis.

**Objective:** FC analysis is predominantly correlational. However, correlation does not necessarily imply causation. This study aims to infer causal functional connectivity (CFC) from functional magnetic resonance imaging (fMRI) data and obtain the sub-networks of CFC that are altered in AD compared to cognitively normal (CN) subjects.

**Methods:** We used the recently developed Time-aware PC algorithm to infer CFC between brain regions. The CFC outcome was compared with correlation-based functional connectivity obtained by sparse partial correlation. Then, Network-based Statistics (NBS) was used to obtain CFC sub-networks that altered in AD subjects compared to healthy controls while correcting for multiple comparisons at 5% level of significance.

**Results:** Our findings identified causal brain networks involving the inferior frontal gyrus, superior temporal gyrus (temporal pole), middle temporal gyrus (temporal pole), and different lobes of the cerebellum to be significantly reduced in strength in AD compared to CN group (p-value = 0.0299; NBS corrected). In the sample dataset that has been analysed, no brain networks were found to exhibit significant increase in strength in AD compared to CN group at 5% level of significance with NBS correction.

**Conclusions:** Our findings provide insights into disruptions in causal brain networks in AD. The corresponding brain regions are in agreement with published medical literature on brain regions impacted by AD. Our work establishes a methodology for finding causal brain networks that are affected by AD using TPC algorithm to compute subject-specific CFC and then using NBS for finding CFC subnetworks that alter between AD and CN groups. Larger datasets are expected to identify further subnetworks affected by AD.

## 1 Introduction

Alzheimer’s disease (AD), a leading cause of dementia worldwide, is characterized by a progressive decline in cognitive function such as impairments to working memory, episodic memory, attention/executive function (de LaCoste and White III, 1993, Christensen et al., 1997). Such cognitive function deficits in AD are believed to be due to abnormality in connectivity between different regions of the brain (Gomez-Isla and Hyman, 1997, Perry and Hodges, 1999, Delbeuck et al., 2003). Evidence from many anatomical and functional studies suggest that AD is a “disconnection syndrome” (Geschwind, 1965, Bozzali et al., 2011, Brier et al., 2014a). Anatomically, AD is characterized by progressive diffuse cortical atrophy, primarily located in the mesial-temporal regions (Yao et al., 2010, Amlien and Fjell, 2014). On the other hand, studies of functional connectivity (FC), which measures the degree of temporal synchrony between the activity of different brain regions, have found evidence of disrupted functional connectivity between brain regions (Wang et al., 2007, Zhao et al., 2020, Biswas and Sripada, 2023).

Several FC analyses measure the degree of statistical association between signals from different brain regions, in other words, associative FC (AFC) (Sheline and Raichle, 2013, Brier et al., 2014b, Briels et al., 2020). However, AFC does not account for the directionality and causality of information flow between regions (Biswas and Shlizerman, 2022a,b). In contrast to AFC, causal functional connectivity (CFC) extends beyond associations and captures the degree of causal interactions between brain regions. Thereby, CFC promises to be more informative than AFC. CFC represents functional connectivity between brain regions by a directed graph, with nodes as the brain regions, directed edges between nodes indicating causal relationships between the brain regions, and weights of the directed edges quantifying the strength of the corresponding causal relationship (Spirtes et al., 2000, Biswas and Shlizerman, 2022a). Yet, there are very few studies that explore causality in functional connectivity (Reid et al., 2019). In this paper, we aim to obtain CFC sub-networks that exhibit significant alteration in AD.

Functional magnetic resonance imaging (fMRI) is a widespread technique to record activity of different brain regions (Logothetis et al., 2001, Smith, 2004, Logothetis, 2008). It measures the Blood Oxygen Level Dependent (BOLD) signal from different brain regions. Studies in functional connectivity have popularly used fMRI data to obtain functional connectivity (Van Den Heuvel and Pol, 2010, Rogers et al., 2007, Kim and Ye, 2020). In an earlier work by Biswas and Sripada (2023), the authors obtained CFC in AD from fMRI data and performed an edge-wise analysis of differences in CFC edges between AD and CN groups. In contrast, in this present paper, we use network-based statistics (NBS) to perform a network-based comparison of the CFC resulting in altered CFC sub-networks in AD compared to CN subjects, while correcting for multiple comparisons in those sub-networks (Zalesky et al., 2010).

In this paper, we use fMRI recordings to obtain CFC for each individual subject. We infer the CFC from fMRI data using the recently developed Time-aware PC algorithm (Biswas and Shlizerman, 2022b). The Time-aware PC algorithm infers causality in neural interactions instead of correlation, in a model-free non-parametric manner, computes whole brain-wide interactions, incorporates multi-regional interactions, and its utility has been demonstrated in fMRI data (Biswas and Sripada, 2023). We then use network-based statistics (NBS) to obtain CFC subnetworks that significantly alter between the subjects with AD and cognitively normal (CN) subjects, while correcting for multiple comparisons (Zalesky et al., 2010, Zhan et al., 2016, 2019). The resulting brain regions are supported by medical literature on brain regions impacted by Alzheimer’s disease.

## 2. Materials and Methods

### 2.1 Participants

The resting fMRI and demographic data were downloaded from the Alzheimer’s Disease Neuroimaging Initiative (ADNI; http://adni.loni.usc.edu/). A total of 129 subjects were included in the study: 41 subjects who are CN, 54 subjects with MCI, and 34 subjects with AD.

Table 1 includes a summary of the participants’ demographic and medical information. In the experiments, the subjects with AD presented significantly lower scores in the screening assessment cognitive test Mini-Mental State Examination (MMSE) in comparison with the other groups. The subjects were age-matched (Kruskal-Wallis test: *p >* 0.8), gender-matched (Chi-Squared test: *p >* 0.1), and matching number of years of education (Kruskal-Wallis test: *p >* 0.2). As expected, MMSE scores had a significant difference between all pairs of groups (Kruskal-Wallis test: *p <* 10^*−*14^).

**Table 1:** Summary of demographic information and Mini Mental State Examination (MMSE) for CN, MCI and AD subjects. The second to fourth columns present group characteristics, mean *±*SD. The fifth column presents *P* values for the statistical significance of the inter-group differences. Differences in Sex was assessed using a Chi-Squared test and differences in Age, Education and MMSE using non-parametric analysis of variance by Kruskal-Wallis test.

| Characteristic | CN | MCI | AD | $P$ |
| --- | --- | --- | --- | --- |
| Number of subjects | 41 | 54 | 34 | - |
| Sex (M/F) | 19/22 | 29/26 | 16/18 | 0.16 |
| Age (years) | $74.9 \pm 6.4$ | $74.2 \pm 7.1$ | $74.4 \pm 7.4$ | 0.86 |
| Education (years) | $16.5 \pm 2.3$ | $15.7 \pm 2.6$ | $15.4 \pm 2.5$ | 0.22 |
| MMSE | $29.1 \pm 1.4$ | $27.8 \pm 1.9$ | $21.9 \pm 4.2$ | $< 10^{-14}$ |

### 2.2 Image Acquisition

The acquisition of fMRI images was performed using Philips Medical Systems scanner. The fMRI images were obtained using an echo planar imaging sequence at a field strength of 3.0 Tesla, with a repetition time (TR) of 3 seconds, an echo time (TE) of 30 milliseconds, and a flip angle of 80 degrees. The matrix size was 64 *×* 64 pixels, 140 volumes, 48 slices per volume, slice thickness of 3.3 mm, and voxel size of 3.3 *×* 3.3 *×* 3.3 mm^3^.

### 2.3 fMRI Preprocessing

The fMRI pre-processing steps were carried out using the CONN toolbox version 21a, which utilizes the Statistical Parametric Mapping (SPM12), both of which are MATLAB-based cross-platform software (Nieto-Castanon, 2021, Friston et al., 1994). We used the default pre-processing pipeline in CONN, consisting of the following steps in order: functional realignment and unwarp (subject motion estimation and correction), functional centering to (0,0,0) coordinates (translation), slice-time correction with interleaved slice order, outlier identification using Artifact Detection and Removal Tool, segmentation into gray matter, white matter and cerebrospinal fluid tissue, and direct normalization into standard Montreal Neurological Institute (MNI) brain space, and lastly, smoothing using spatial convolution with a Gaussian kernel of 8mm full-width half maximum. This pipeline was followed by detrending and bandpass filtering (0.001-0.1 Hz) to remove low-frequency scanner drift and physiological noise in the fMRI images. The first four time points have been filtered out to remove any artifacts.

For the extraction of Regions-Of-Interest (ROIs), the automated anatomical labeling (AAL) atlas was utilized on the pre-processed rs-fMRI dataset (Tzourio-Mazoyer et al., 2002). The list of all regions in the AAL atlas is provided in Appendix Appendix A along with their abbreviated, short, and full region names. This parcellation method has been demonstrated to be optimal for studying the FC between brain regions (Arslan et al., 2018). The voxels within each ROI were averaged, resulting in a time series for each ROI.

### 2.4 Functional Connectivity Analyses

#### 2.4.1 Associative FC

We employed Sparse Partial Correlation to obtain AFC from fMRI time series data (Banerjee et al., 2008, Pervaiz et al., 2020, Schmittmann et al., 2015). The sparse partial correlation regularizes partial correlation with a graphical LASSO penalty, whose regularization parameters were selected by cross-validation (Meinshausen et al., 2006).

#### 2.4.2 Causal FC

In our study, CFC is inferred using the Time-aware PC (TPC) Algorithm, a recent method to infer CFC from brain signals (Biswas and Shlizerman, 2022b, Biswas and Mukherjee, 2024). By implementing the Directed Markov Property in a time series setting, the TPC algorithm models causal interactions as an unrolled Directed Acyclic Graph (DAG), representing the time series’ spatial and temporal interactions. In this unrolled DAG, nodes represent regions of interest at specific times, and edges indicate directional causal influences between these nodes. The TPC algorithm transforms time series data into sequential variables with a maximum time delay (*τ* = 1), and applies the Peter-Clark (PC) algorithm for DAG estimation, later rolling the DAG back to form a comprehensive CFC graph. This method allows for the analysis of complex, multivariate interactions and supports whole-brain CFC estimation, accommodating the BOLD signal’s temporal delays and feedback loops (Biswas and Sripada, 2023). The Python package *Time-Aware PC* is used for implementing this approach. Our methodology is particularly suited to fMRI data, accounting for the relatively slow temporal resolution of the BOLD signal and enabling the detection of both sequential and contemporaneous interactions within the brain network (Biswas and Sripada, 2023). The outcome of this approach offers a non-parametric, detailed mapping of causal functional connectivity across the brain, enhancing our understanding of the directional flow of information and its disruption in AD.

### 2.5 Alterations of CFC edges in Alzheimer’s disease

NBS is a popular non-parametric statistical method for identifying inter-group differences in large brain networks while controlling for multiple comparisons (Zalesky et al., 2010, Zhan et al., 2016, Olivito et al., 2017, De Schipper et al., 2018, Zhan et al., 2019, Zhu et al., 2023). It is a method for controlling the FWER when a large number of univariate tests are conducted over the connections in the network. NBS method outputs the connected components in the brain network that exhibit inter-group difference and a corresponding FWER-corrected p-value for each such component. The FWER-corrected p-values are calculated for each component using permutation testing. We used 10,000 random permutations for the permutation tests. For implementation, we used the *NBS Matlab Toolbox* and its extension for directed networks (https://www.nitrc.org/projects/nbs/) (Zalesky et al., 2010). In the implementation of NBS, we used a primary threshold of 1.8 and FWER corrected significance level of 0.05. At that significance level, the critical threshold of cluster size was 6. For finding altered sub-networks between CN and AD groups, NBS considers the direction of each connection in the whole CFC among subjects in CN and AD groups. We outlined detailed steps of implementing NBS in Appendix B.

We applied NBS to the subject-specific CFCs estimated by TPC, to identify connected components of the CFC that are altered in strength for subjects with AD compared to the CN.

## 3. Results

### 3.1 Subject-specific CFC

Figure 1 shows the AFC obtained by sparse partial correlation and CFC obtained by TPC for an example subject who is CN. The AFC (top row) revealed an extensive associative network, as evidenced by dense symmetric adjacency matrix and dense undirected graphical representations. It is expected that the CFC will be a directed subgraph of the AFC and be consistent with the overall patterns present in the AFC. The overall patterns present in the CFC obtained by TPC indeed match with the AFC as visualized in their adjacency matrices (See Figure 1-left column). On a detailed level, there are differences between TPC-CFC and AFC: TPC results in a directed graph thereby having an asymmetric adjacency matrix while AFC is an undirected graph with symmetric adjacency matrix. Furthermore, in contrast to the dense adjacency matrix and graphical representations of the AFC, the CFC obtained by TPC results in a sparse adjacency matrix and sparse graphical representation devoid of noise since the connections are filtered by conditional dependence tests in the TPC algorithm.

**Figure 1.**
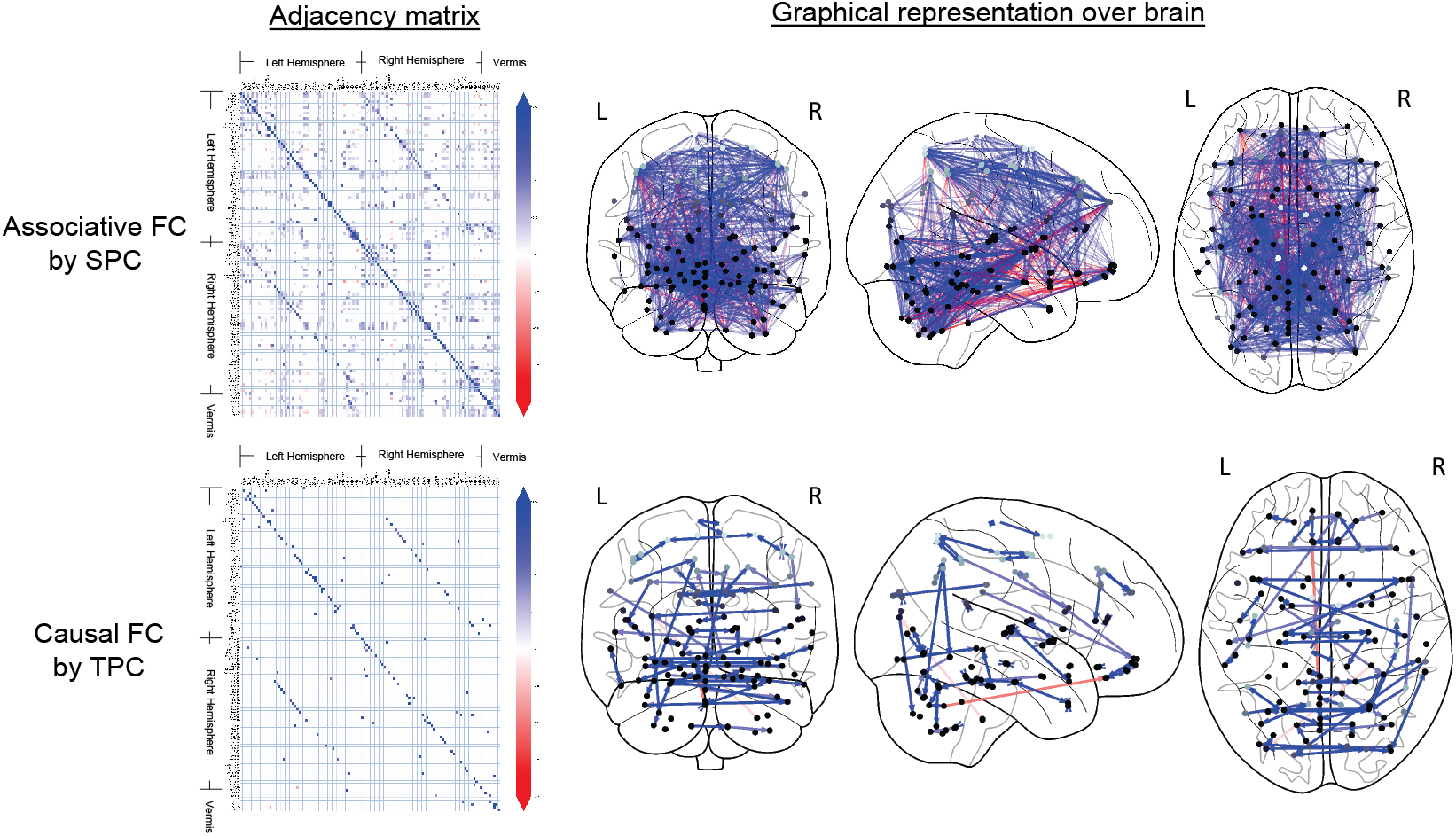
CFC for an example subject who is CN, estimated by TPC algorithm. The estimated CFC is represented by its adjacency matrix (left column), whose non-zero entry (*i, j*) represents the connection of region *i→ j*. The CFC is also visualized with directed graph edges on the Frontal, Axial and Lateral brain maps (2nd to 4th columns from left). The nodes correspond to brain region centers, ranging from superficial (light gray) to deeper (darker gray) regions, in the AAL brain atlas. The brain regions are annotated by Left (L) and Right (R) hemispheres of the brain and Vermis (V).

### 3.2 Altered CFC sub-networks in AD

Figure 2 shows the significantly altered (*P <* 0.05; NBS corrected) CFC sub-networks in AD compared to CN. In the implementation of NBS, t-statistic thresholds of 0.1 to 3.1 in steps of 0.1 were tried out. Among those, thresholds of 1.8, 2.2 and 2.3 yielded altered sub-networks at 5% level of significance. At each of these thresholds, a sparse thresholded adjacency matrix is obtained (Figure 2-left row). A subset of the thresholded edges is outputted by NBS as an altered sub-network at 5% level of significance. Figure 2 shows the altered sub-network’s adjacency matrix representation (2nd column from left) as well as graphical representation (3rd to 5th columns from left) on brain schematics.

**Figure 2.**
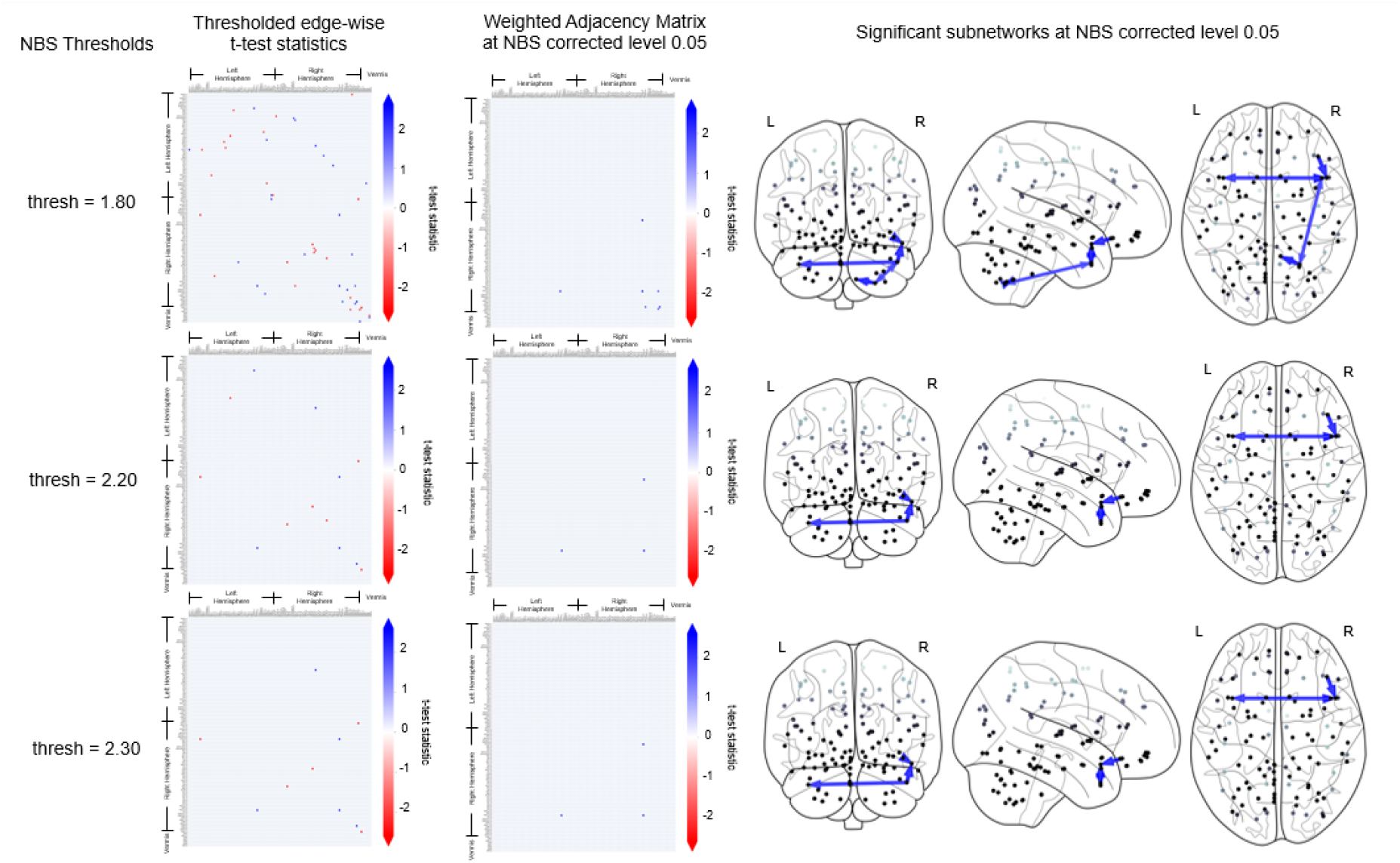
This figures shows the significantly altered sub-networks obtained by NBS implemented on the CFC outcome of TPC. For NBS test-statistics thresholds of 1.8, 2.2 and 2.3 (top to bottom), the thresholded edge-wise t-test statistics (left column) and the result of the NBS corrected sub-network are shown: weighted adjacency matrix representation (2nd column from left), graphical representation on brain schematics on the Frontal, Axial and Lateral brain maps (3rd to 5th columns from left).

### 3.3 Brain regions in the altered CFC sub-networks

In Table 2, we report the CFC edges that are present in the altered CFC sub-networks with greater average strength in CN subjects compared to AD at NBS corrected 5% level of significance, and their source brain regions. The corresponding brain regions are in agreement with published medical literature cited in Table 2. No significantly altered sub-networks were obtained by NBS for lower average strength in CN subjects compared to AD at 5% level of significance.

**Table 2:**
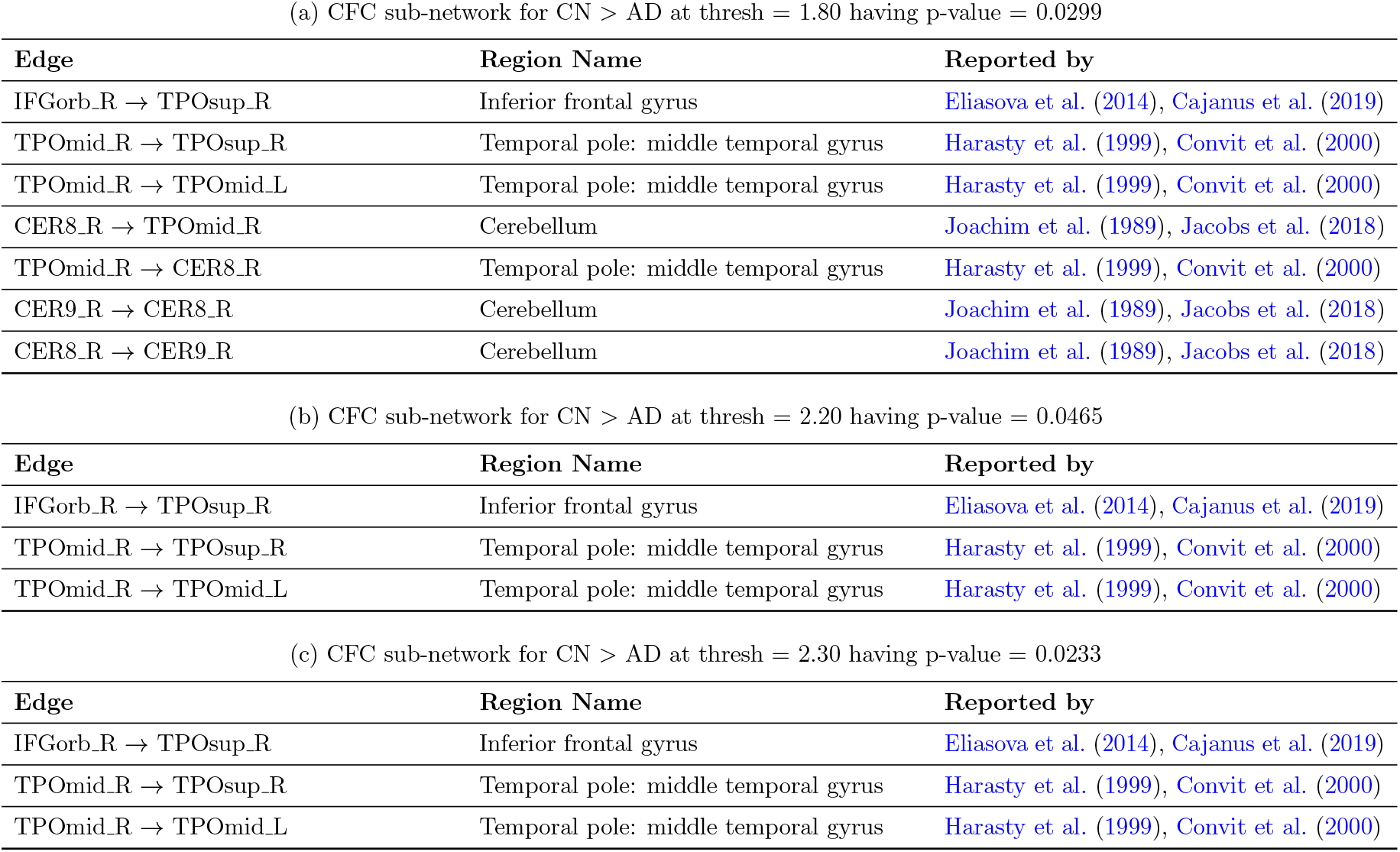
CFC edges in significantly altered CFC sub-networks with greater weight in CN compared to AD group at NBS corrected 5% level of significance. The sub-networks are obtained at different NBS’s t-statistic thresholds: (a) 1.80, (b) 2.20 and (c) 2.30 yielding altered sub-networks with p-values of 0.0299, 0.0465 and 0.0233 respectively. The corresponding source brain regions are in agreement with regions reported in literature (right column) as impacted by AD.

## 4. Discussion

In this study, we obtained CFC sub-networks of the whole brain that are altered in AD compared to CN from resting state fMRI time series. We used the recently developed TPC algorithm based on directed graphical modeling in time series, to compute the CFC. We first computed the subject-specific CFC using TPC and compared it with AFC obtained by sparse partial correlation. We then used the CFC outcomes of TPC for further investigation into altered CFC sub-networks in AD. In this regard, we used NBS to obtain CFC sub-networks that alter in AD compared to CN subjects at 5% level of significance, while correcting for multiple comparisons in those sub-networks. The brain regions in those altered sub-networks were found to be in agreement with medical literature for regions impacted by AD.

In Figure 2 and Table 2, the presence of CFC sub-network with weight of edges in AD greater than that in CN is consistent with published studies in the literature. The absence of CFC sub-networks with weight of edges in AD lower than that in CN at 5% level of significance, supports the hypothesis that AD is a “disconnection syndrome” (Geschwind, 1965, Bozzali et al., 2011, Brier et al., 2014a).

In Table 2, the brain regions of Inferior frontal gyrus (Inferior frontal gyrus, pars orbitalis Right → Temporal pole: superior temporal gyrus Right) and Temporal pole: middle temporal gyrus (Temporal pole: middle temporal gyrus Right → Temporal pole: superior temporal gyrus Right, → Temporal pole: middle temporal gyrus Right → Temporal pole: middle temporal gyrus Left) are prominent in sub-networks with lower CFC weight in AD compared to CN subjects. Among these regions, the Inferior frontal gyrus plays a role in attentional control (Hampshire et al., 2010), and language processing (Rota et al., 2009), and is known to be impacted by AD (Eliasova et al., 2014, Cajanus et al., 2019). The temporal pole: middle temporal gyrus, another region in Table 2, is considered to be important in word memory and meaning (Luria, 1976, Benson, 1980, Ojemann and Mateer, 1979, Damasio and Damasio, 1980, Frackowiak et al., 1981, Mesulam, 1990, Demonet et al., 1993, Gazzaniga, 1993). It has been reported to be impacted by AD (Harasty et al., 1999, Convit et al., 2000). Also present in Table 2(a) is the Cerebellum (Lobule VIII of cerebellar hemisphere Right *→* Temporal pole: middle temporal gyrus Right, Lobule IX of cerebellar hemisphere Right *→* Lobule VIII of cerebellar hemisphere Right, Lobule VIII of cerebellar hemisphere Right *→* Lobule IX of cerebellar hemisphere Right), which plays an essential role in motor control and perception (Paulin, 1993, Akshoomoff and Courchesne, 1992, Botez et al., 1989), and has been reported to be impacted by AD (Joachim et al., 1989, Jacobs et al., 2018).

### 4.1 Conclusion

In this paper, we have demonstrated the following: (a) Application of the TPC algorithm to compute whole-brain CFC for each subject (b) Comparison of CFC with AFC computed using sparse partial correlation (c) Identified CFC sub-networks with altered connectivity in subjects with AD compared to CN using NBS at 5% level of significance. (d) Interpretation of altered CFC sub-networks in the context of AD using domain (neuropathological) knowledge. The findings are consistent with published medical literature. The results demonstrate the potential of computing the whole-brain CFC from fMRI data using the TPC algorithm to gain prognostic and diagnostic insights.

## Appendix A. Automated Anatomical Labeling (AAL) Atlas

The regions in the AAL atlas along with their abbreviated, short and full names are listed in Table A.3.

**Table A.3:**
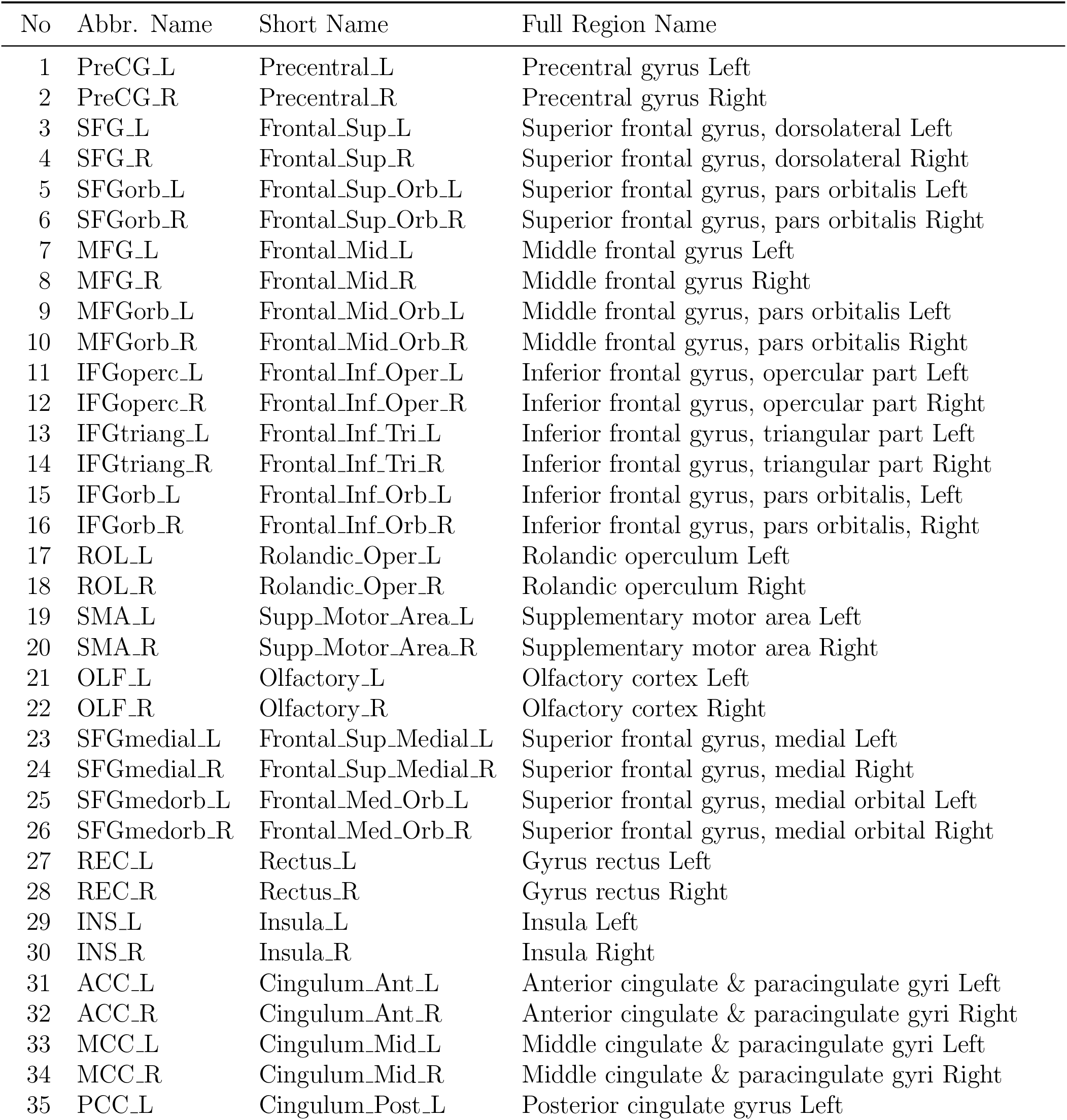
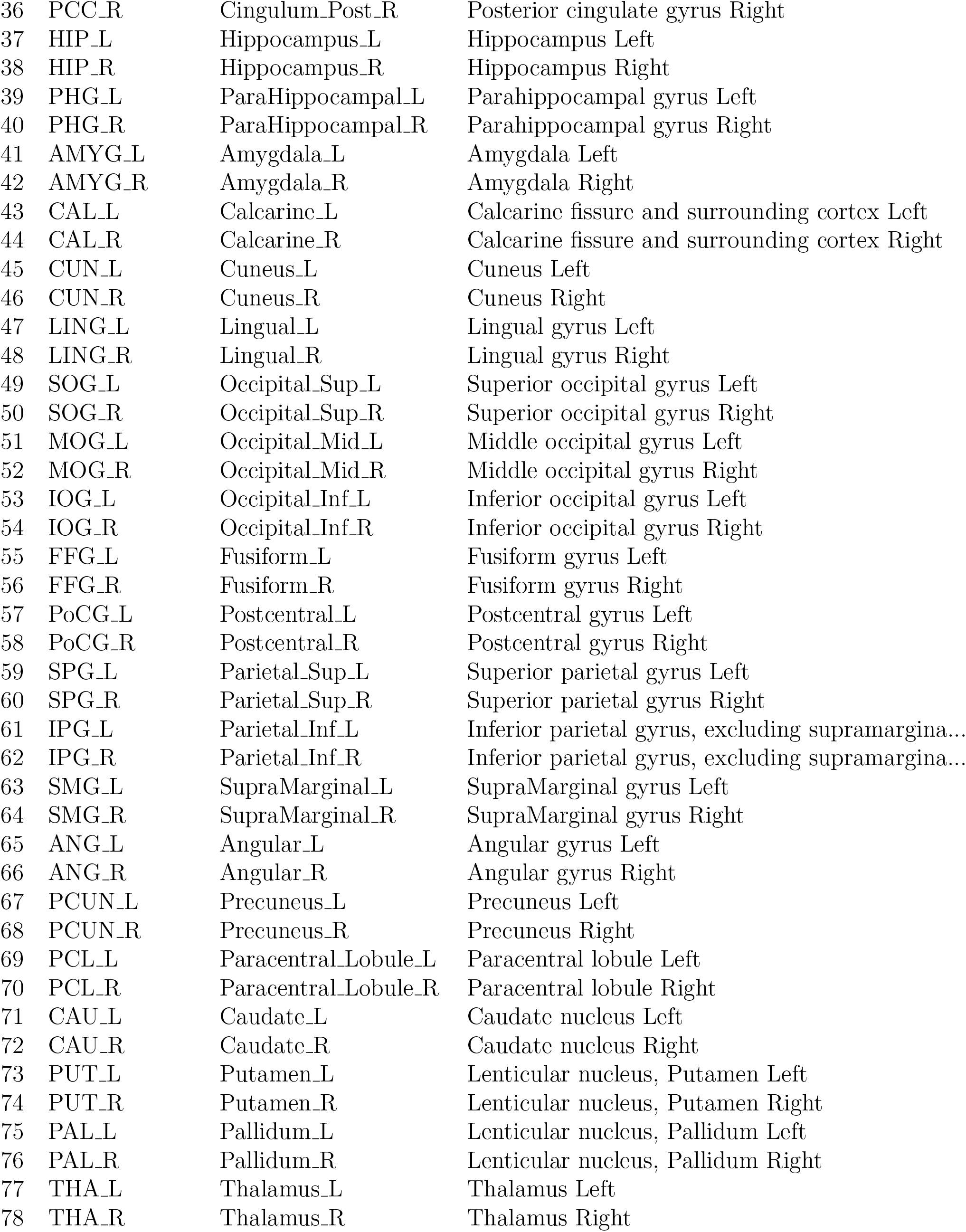
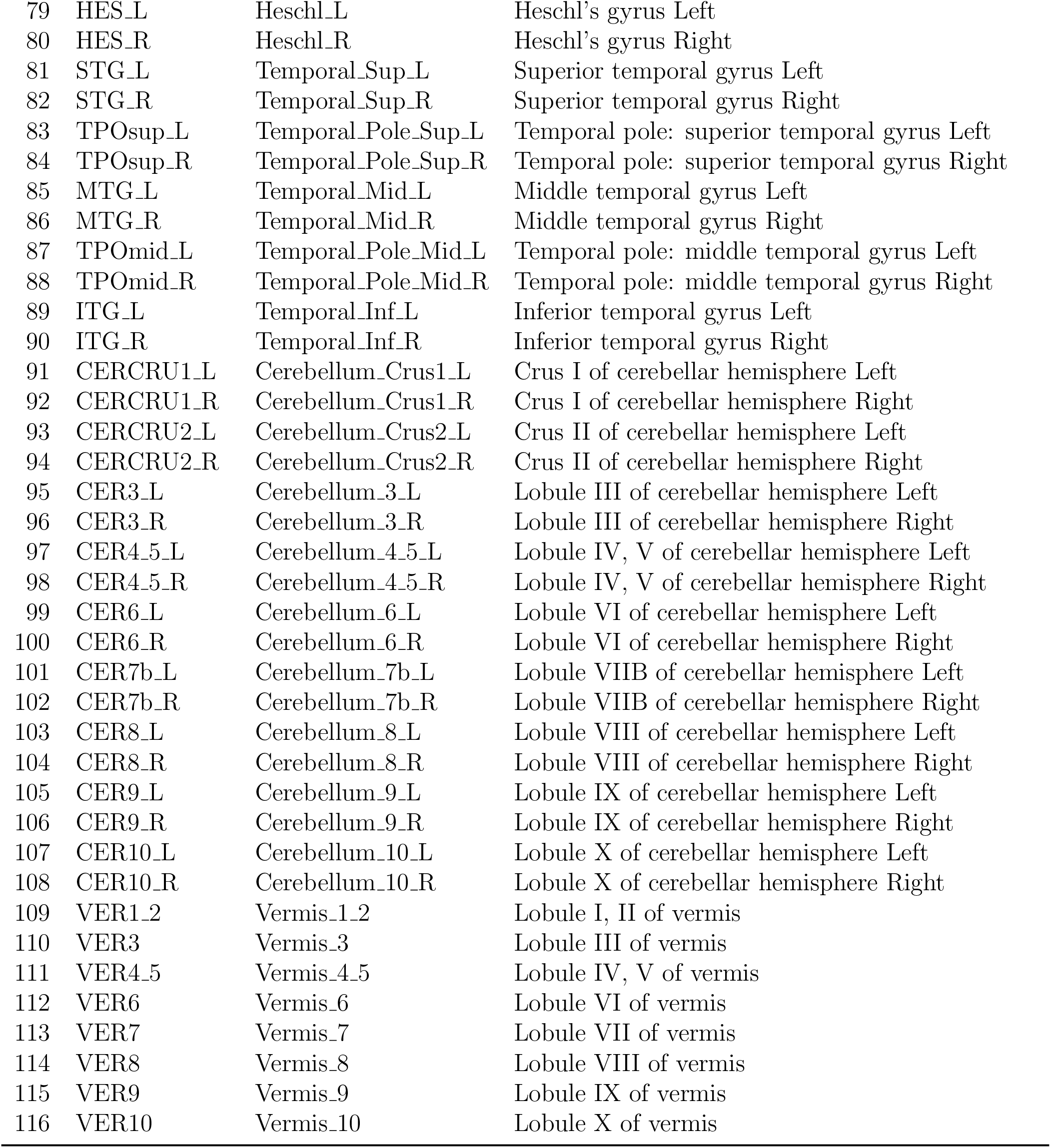
Names of regions in the AAL Atlas.

## Appendix B. The Network-Based Statistics (NBS) method

For comparing the CFCs between two groups, NBS initially calculates a Generalized Linear Model based statistic to score each connection. These scores are then subjected to a primary threshold, leading to the selection of suprathreshold links with scores surpassing this threshold. The thresholding is done separately for positive and negative scores, identifying connected components where individuals with CN exhibit notably higher or lower connectivity strength than those with AD. Various primary thresholds were evaluated in the range of 0.1 to 3 in steps of 0.1, and fixed at 1.8 which is the lowest in the range yielding significance at family wise error rate (FWER) level 0.05. Although the primary threshold impacts method sensitivity, yet FWER control is ensured regardless of the chosen threshold (Zalesky et al., 2010). Subsequently, components within the suprathreshold link set are identified, and their sizes are recorded. To establish component significance, a non-parametric permutation test is conducted. In each permutation, subjects are randomly reassigned to CN and AD groups. Independent scores are computed for each link, and the size of the largest connected component among suprathreshold links is noted. This process is reiterated 10,000 times to generate a null distribution of the pseudo size of the connected component. Ultimately, the corrected p-value for a connected component of size M is determined based on the proportion of permutations where the largest connected component size exceeds M. The NBS method is implemented in the *NBS Matlab Toolbox* and its extension for directed networks (https://www.nitrc.org/projects/nbs/) (Zalesky et al., 2010).

## Notes

### Competing Interest Statement

The authors have declared no competing interest.

### Summary of Updates

Updated a typographical error in the name of affiliation of second author: replaced "&" with "and"

## References

Akshoomoff, N. A. and Courchesne, E. (1992). A new role for the cerebellum in cognitive operations. Behavioral neuroscience, 106(5):731.

Amlien, I. and Fjell, A. (2014). Diffusion tensor imaging of white matter degeneration in alzheimer’s disease and mild cognitive impairment. Neuroscience, 276:206–215.

Arslan, S., Ktena, S. I., Makropoulos, A., Robinson, E. C., Rueckert, D., and Parisot, S. (2018). Human brain mapping: A systematic comparison of parcellation methods for the human cerebral cortex. Neuroimage, 170:5–30.

Banerjee, O., Ghaoui, L. E., and d’Aspremont, A. (2008). Model selection through sparse maximum likelihood estimation for multivariate gaussian or binary data. Journal of Machine learning research, 9(Mar):485–516.

Benson, D. F. (1980). Aphasia, alexia, and agraphia. Archives of Neurology.

Biswas, R. and Mukherjee, S. (2024). Consistent causal inference from time series with pc algorithm and its time-aware extension. Statistics and Computing, 34(1):14.

Biswas, R. and Shlizerman, E. (2022a). Statistical perspective on functional and causal neural connectomics: A comparative study. Frontiers in Systems Neuroscience, 16.

Biswas, R. and Shlizerman, E. (2022b). Statistical perspective on functional and causal neural connectomics: The time-aware pc algorithm. PLOS Computational Biology, 18(11):1–27.

Biswas, R. and Sripada, S. (2023). Causal functional connectivity in alzheimer’s disease computed from time series fmri data. Frontiers in Computational Neuroscience, 17.

Botez, M. I., Botez, T., Elie, R., and Attig, E. (1989). Role of the cerebellum in complex human behavior. The Italian Journal of Neurological Sciences, 10:291–300.

Bozzali, M., Padovani, A., Caltagirone, C., and Borroni, B. (2011). Regional grey matter loss and brain disconnection across alzheimer disease evolution. Current medicinal chemistry, 18(16):2452–2458.

Briels, C. T., Schoonhoven, D. N., Stam, C. J., de Waal, H., Scheltens, P., and Gouw, A. A. (2020). Reproducibility of eeg functional connectivity in alzheimer’s disease. Alzheimer’s research & therapy, 12:1–14.

Brier, M. R., Thomas, J. B., and Ances, B. M. (2014a). Network dysfunction in alzheimer’s disease: refining the disconnection hypothesis. Brain connectivity, 4(5):299– 311.

Brier, M. R., Thomas, J. B., Fagan, A. M., Hassenstab, J., Holtzman, D. M., Benzinger, T. L., Morris, J. C., and Ances, B. M. (2014b). Functional connectivity and graph theory in preclinical alzheimer’s disease. Neurobiology of aging, 35(4):757–768.

Cajanus, A., Solje, E., Koikkalainen, J., Lötjönen, J., Suhonen, N.-M., Hallikainen, I., Vanninen, R., Hartikainen, P., de Marco, M., Venneri, A., et al. (2019). The association between distinct frontal brain volumes and behavioral symptoms in mild cognitive impairment, alzheimer’s disease, and frontotemporal dementia. Frontiers in neurology, 10:1059.

Christensen, H., Griffiths, K., MacKinnon, A., and Jacomb, P. (1997). A quantitative review of cognitive deficits in depression and alzheimer-type dementia. Journal of the International Neuropsychological Society, 3(6):631–651.

Convit, A., De Asis, J., De Leon, M., Tarshish, C., De Santi, S., and Rusinek, H. (2000). Atrophy of the medial occipitotemporal, inferior, and middle temporal gyri in non-demented elderly predict decline to alzheimer’s disease. Neurobiology of aging, 21(1):19–26.

Damasio, H. and Damasio, A. R. (1980). The anatomical basis of conduction aphasia. Brain, 103(2):337–350.

de LaCoste, M.-C. and White III, C. L. (1993). The role of cortical connectivity in alzheimer’s disease pathogenesis: a review and model system. Neurobiology of aging, 14(1):1–16.

De Schipper, L. J., Hafkemeijer, A., Van der Grond, J., Marinus, J., Henselmans, J. M., and Van Hilten, J. J. (2018). Altered whole-brain and network-based functional connectivity in parkinson’s disease. Frontiers in neurology, 9:419.

Delbeuck, X., Van der Linden, M., and Collette, F. (2003). Alzheimer’disease as a disconnection syndrome? Neuropsychology review, 13:79–92.

Demonet, J., Wise, R., and Frackowiak, R. (1993). Language functions explored in normal subjects by positron emission tomography: A critical review. Human Brain Mapping, 1(1):39–47.

Eliasova, I., Anderkova, L., Marecek, R., and Rektorova, I. (2014). Non-invasive brain stimulation of the right inferior frontal gyrus may improve attention in early alzheimer’s disease: a pilot study. Journal of the neurological sciences, 346(1-2):318–322.

Frackowiak, R., Pozzilli, C., Legg, N. d., Du Boulay, G., Marshall, J., Lenzi, G. L., and Jones, T. (1981). Regional cerebral oxygen supply and utilization in dementia. a clinical and physiological study with oxygen-15 and positron tomography. Brain: a journal of neurology, 104(Pt 4):753–778.

Friston, K. J., Holmes, A. P., Worsley, K. J., Poline, J.-P., Frith, C. D., and Frackowiak, R. S. (1994). Statistical parametric maps in functional imaging: a general linear approach. Human brain mapping, 2(4):189–210.

Gazzaniga, M. S. (1993). Language and the cerebral hemispheres. Discussions in Neuroscience, 10(1):106–108.

Geschwind, N. (1965). Disconnexion syndromes in animals and man. Brain, 88(3):585– 585.

Gomez-Isla, T. and Hyman, B. (1997). Connections and cognitive impairment in alzheimer’s disease. In Connections, cognition and alzheimer’s disease, pages 149–166. Springer.

Hampshire, A., Chamberlain, S. R., Monti, M. M., Duncan, J., and Owen, A. M. (2010). The role of the right inferior frontal gyrus: inhibition and attentional control. Neuroimage, 50(3):1313–1319.

Harasty, J. A., Halliday, G. M., Kril, J., and Code, C. (1999). Specific temporoparietal gyral atrophy reflects the pattern of language dissolution in alzheimer’s disease. Brain, 122(4):675–686.

Jacobs, H. I., Hopkins, D. A., Mayrhofer, H. C., Bruner, E., van Leeuwen, F. W., Raaijmakers, W., and Schmahmann, J. D. (2018). The cerebellum in alzheimer’s disease: evaluating its role in cognitive decline. Brain, 141(1):37–47.

Joachim, C. L., Morris, J. H., and Selkoe, D. J. (1989). Diffuse senile plaques occur commonly in the cerebellum in alzheimer’s disease. The American journal of pathology, 135(2):309.

Kim, B.-H. and Ye, J. C. (2020). Understanding graph isomorphism network for rs-fmri functional connectivity analysis. Frontiers in neuroscience, 14:545464.

Logothetis, N. K. (2008). What we can do and what we cannot do with fmri. Nature, 453(7197):869–878.

Logothetis, N. K., Pauls, J., Augath, M., Trinath, T., and Oeltermann, A. (2001). Neurophysiological investigation of the basis of the fmri signal. nature, 412(6843):150– 157.

Luria, A. R. (1976). The working brain: An introduction to neuropsychology.

Meinshausen, N., Bühlmann, P., et al. (2006). High-dimensional graphs and variable selection with the lasso. The annals of statistics, 34(3):1436–1462.

Mesulam, M.-M. (1990). Large-scale neurocognitive networks and distributed processing for attention, language, and memory. Annals of Neurology: Official Journal of the American Neurological Association and the Child Neurology Society, 28(5):597–613.

Nieto-Castanon, A. (2021). Conn functional connectivity toolbox (rrid: Scr 009550), version 21.

Ojemann, G. and Mateer, C. (1979). Human language cortex: localization of memory, syntax, and sequential motor-phoneme identification systems. Science, 205(4413):1401– 1403.

Olivito, G., Cercignani, M., Lupo, M., Iacobacci, C., Clausi, S., Romano, S., Masciullo, M., Molinari, M., Bozzali, M., and Leggio, M. (2017). Neural substrates of motor and cognitive dysfunctions in sca2 patients: a network based statistics analysis. NeuroImage: Clinical, 14:719–725.

Paulin, M. G. (1993). The role of the cerebellum in motor control and perception. Brain, behavior and evolution, 41(1):39–50.

Perry, R. J. and Hodges, J. R. (1999). Attention and executive deficits in alzheimer’s disease: A critical review. Brain, 122(3):383–404.

Pervaiz, U., Vidaurre, D., Woolrich, M. W., and Smith, S. M. (2020). Optimising network modelling methods for fmri. Neuroimage, 211:116604.

Reid, A. T., Headley, D. B., Mill, R. D., Sanchez-Romero, R., Uddin, L. Q., Marinazzo, D., Lurie, D. J., Valdés-Sosa, P. A., Hanson, S. J., Biswal, B. B., et al. (2019). Advancing functional connectivity research from association to causation. Nature neuroscience, 1(10).

Rogers, B. P., Morgan, V. L., Newton, A. T., and Gore, J. C. (2007). Assessing functional connectivity in the human brain by fmri. Magnetic resonance imaging, 25(10):1347–1357.

Rota, G., Sitaram, R., Veit, R., Erb, M., Weiskopf, N., Dogil, G., and Birbaumer, N. (2009). Self-regulation of regional cortical activity using real-time fmri: The right inferior frontal gyrus and linguistic processing. Human brain mapping, 30(5):1605– 1614.

Schmittmann, V. D., Jahfari, S., Borsboom, D., Savi, A. O., and Waldorp, L. J. (2015). Making large-scale networks from fmri data. PloS one, 10(9):e0129074.

Sheline, Y. I. and Raichle, M. E. (2013). Resting state functional connectivity in preclinical alzheimer’s disease. Biological psychiatry, 74(5):340–347.

Smith, S. M. (2004). Overview of fmri analysis. The British Journal of Radiology, 77(Suppl 2):S167–S175.

Spirtes, P., Glymour, C. N., Scheines, R., and Heckerman, D. (2000). Causation, prediction, and search. MIT press.

Tzourio-Mazoyer, N., Landeau, B., Papathanassiou, D., Crivello, F., Etard, O., Delcroix, N., Mazoyer, B., and Joliot, M. (2002). Automated anatomical labeling of activations in spm using a macroscopic anatomical parcellation of the mni mri single-subject brain. Neuroimage, 15(1):273–289.

Van Den Heuvel, M. P. and Pol, H. E. H. (2010). Exploring the brain network: a review on resting-state fmri functional connectivity. European neuropsychopharmacology, 20(8):519–534.

Wang, K., Liang, M., Wang, L., Tian, L., Zhang, X., Li, K., and Jiang, T. (2007). Altered functional connectivity in early alzheimer’s disease: A resting-state fmri study. Human brain mapping, 28(10):967–978.

Yao, Z., Zhang, Y., Lin, L., Zhou, Y., Xu, C., Jiang, T., and Initiative, A. D. N. (2010). Abnormal cortical networks in mild cognitive impairment and alzheimer’s disease. PLoS computational biology, 6(11):e1001006.

Zalesky, A., Fornito, A., and Bullmore, E. T. (2010). Network-based statistic: identifying differences in brain networks. Neuroimage, 53(4):1197–1207.

Zhan, C., Chen, H.-J., Gao, Y.-Q., and Zou, T.-X. (2019). Functional network-based statistics reveal abnormal resting-state functional connectivity in minimal hepatic encephalopathy. Frontiers in Neurology, 10:33.

Zhan, Y., Yao, H., Wang, P., Zhou, B., Zhang, Z., An, N., Ma, J., Zhang, X., Liu, Y., et al. (2016). Network-based statistic show aberrant functional connectivity in alzheimer’s disease. IEEE Journal of Selected Topics in Signal Processing, 10(7):1182–1188.

Zhao, J., Du, Y.-H., Ding, X.-T., Wang, X.-H., and Men, G.-Z. (2020). Alteration of functional connectivity in patients with alzheimer’s disease revealed by resting-state functional magnetic resonance imaging. Neural Regeneration Research, 15(2):285–292.

Zhu, L., Dang, G., Wu, W., Zhou, J., Shi, X., Su, X., Ren, H., Pei, Z., Lan, X., Lian, C., et al. (2023). Functional connectivity changes are correlated with sleep improvement in chronic insomnia patients after rtms treatment. Frontiers in Neuroscience, 17:1135995.

